# The Analgesic and Dissociative Properties of Ketamine are Separate and Correspond to Distinct Neural Mechanisms

**DOI:** 10.1101/2025.07.25.666594

**Authors:** Noam Goldway, Talma Hendler, Itamar Jalon, Yotam Pasternak, Roy Sar-El, Dan Mirelman, Noam Sarna, Nili Green, Yara Agbaria, Haggai Sharon

## Abstract

Ketamine, a psychoactive medication, exerts both analgesic and dissociative effects. However, whether its analgesic effect stems from its dissociative properties is a topic of debate. Our study aimed to determine whether ketamine’s analgesic and dissociative effects are supported by distinct neural mechanisms. In a within-subject, placebo-controlled study, 37 healthy volunteers were administered ketamine (0.4 mg/kg bolus followed by a continuous drip of 0.4 mg/kg/h) or saline during fMRI sessions where thermal pain was induced. Our results indicate that while ketamine significantly reduced thermal pain ratings and produced robust dissociative effects, these outcomes were not correlated. Neurally, ketamine reduced pain-related brain activations across a network of regions, including the insula and anterior cingulate cortex. Additionally, ketamine significantly diminished functional connectivity between default mode network regions, and this reduction was correlated with the intensity of dissociation. These findings suggest that ketamine’s analgesic and dissociative effects are independent and mediated by distinct neural pathways.

## Introduction

In recent years, there has been a renewed interest in the potential medical applications of mind-altering drugs in the treatment of a variety of clinical neuropsychiatric disorders, such as major depression, post traumatic stress disorder (PTSD) and chronic pain^1–4^. As the mechanisms underlying the therapeutic effects of psychedelics have come under renewed investigation, a key question has emerged: to what extent are their clinical benefits dependent on the altered states of consciousness they induce? While some regard the subjective experience as an epiphenomenon of neurochemical action, others argue it is integral to therapeutic efficacy^5–7^.

In this context, an interesting case is the relationship between dissociation and analgesia. Dissociation is a disruption or discontinuity in the normal functioning of cognitive modules such as affect, memory, identity, perception, and motor control^8^. Dissociation is often measured along three dimensions: detachment from self (depersonalization), detachment from surroundings (derealization), and discontinuation of ongoing experience, along with sensory distortions^9^. Clinically, dissociative states were shown to be related to pain hyposensitivity in borderline personality disorder (BPD), and in PTSD^10–13^. Pharmacologically, both dissociation and analgesia can be induced using ketamine^14–16^. The co-occurrence of dissociation and reduced pain sensitivity in both clinical and pharmacological contexts has led some to speculate about a mechanistic link between the two phenomena. Such claims rest on the understanding that pain is a complex experience involving multiple cognitive processes. Some of these processes relate to the objective properties of the pain-inducing stimulus, such as its location, temperature or pressure. Meanwhile, the affective aspects of the pain experience, which involve attention allocation and behavioral modifications such as avoidance or coping behaviors, are associated with the subjective experience of suffering^17–19^. The dissociative state, wherein an individual feels disconnected from their own body and affective state, can potentially modify the threatening, unpleasant nature of the pain experience. This modification can lead to an analgesic effect where the pain is perceived as less intense^20^. An alternate perspective suggests that the analgesic and dissociative effects are independent, stemming from different mechanisms. This debate is fueled by mixed results from past research focusing on ketamine interventions. Support for the notion that the two effects are correlated can be found in clinical guidelines^21^ and empirical findings demonstrating that similar plasma concentrations of ketamine and its metabolites are needed to produce its analgesic and mind-altering effects^22^. However, other studies suggest that ketamine’s analgesic and dissociative effects are not correlated^23,24^. While the diverging conclusions of these studies might be accounted for by different dosing of ketamine given to participants or by the types of statistical models implemented, the question of whether dissociative and analgesic effects of ketamine have distinct mechanisms remains open. A potential approach to address this question could involve examining ketamine’s impact on the neural systems that underlie its analgesic and dissociative effects.

The brain mechanisms supporting acute pain are well-characterized and include activation across a broad network of brain regions such as the insula, dorsal Anterior Cingulate Cortex (dACC), primary (S1) and secondary (S2) somatosensory cortices, and subcortical regions including the thalamus, basal ganglia, amygdala, and periaqueductal gray (PAG)^17^. Early research examining the neural effects of ketamine on the activation of these regions during pain perception indicates that the drug reduces activation in several areas crucial for pain processing, such as the S2, insula, and dACC^25^.

The neural mechanisms underlying ketamine-induced dissociation, however, remain somewhat elusive. Most imaging studies examining the effects of ketamine on brain dynamics during dissociative experiences in healthy individuals utilized functional connectivity measurements during resting state^26–29^. Although findings vary, there has been consistent observation that ketamine leads to the disintegration of the Default Mode Network (DMN) and Salience Network (SN)^27–30^

In the current study, our aim was to determine whether the analgesic and dissociative effects of ketamine are mediated by the same or distinct neural mechanisms. A total of thirty-seven healthy volunteers (mean age 29.7 ± 3.21 years, 21 females) were enrolled in the study, with 33 completing the full procedure (mean age 30.1 ± 3.09 years, 18 females). Each volunteer participated in two sessions: one involving the administration of a placebo (IV saline) and the other involving the administration of racemic ketamine (initial bolus of .4 mg/kg IV over 10 minutes, followed by a continuous drip of .4 mg/kg/h). Before administering either substance, participants underwent a pain calibration procedure (see methods). Approximately ten minutes after the conclusion of the initial bolus infusion, participants responded to a verbally administered Clinician-Administered Dissociative States Scale (CADSS) to assess their dissociative state^9^ (Figure 1.A). Afterward, participants took part in an fMRI session that included a ‘pain task’. During this task, they received either a painful or non-painful thermal stimulus applied to their right leg, determined by the calibration procedure conducted earlier that day (Figure 1.B).

**Figure 1.**
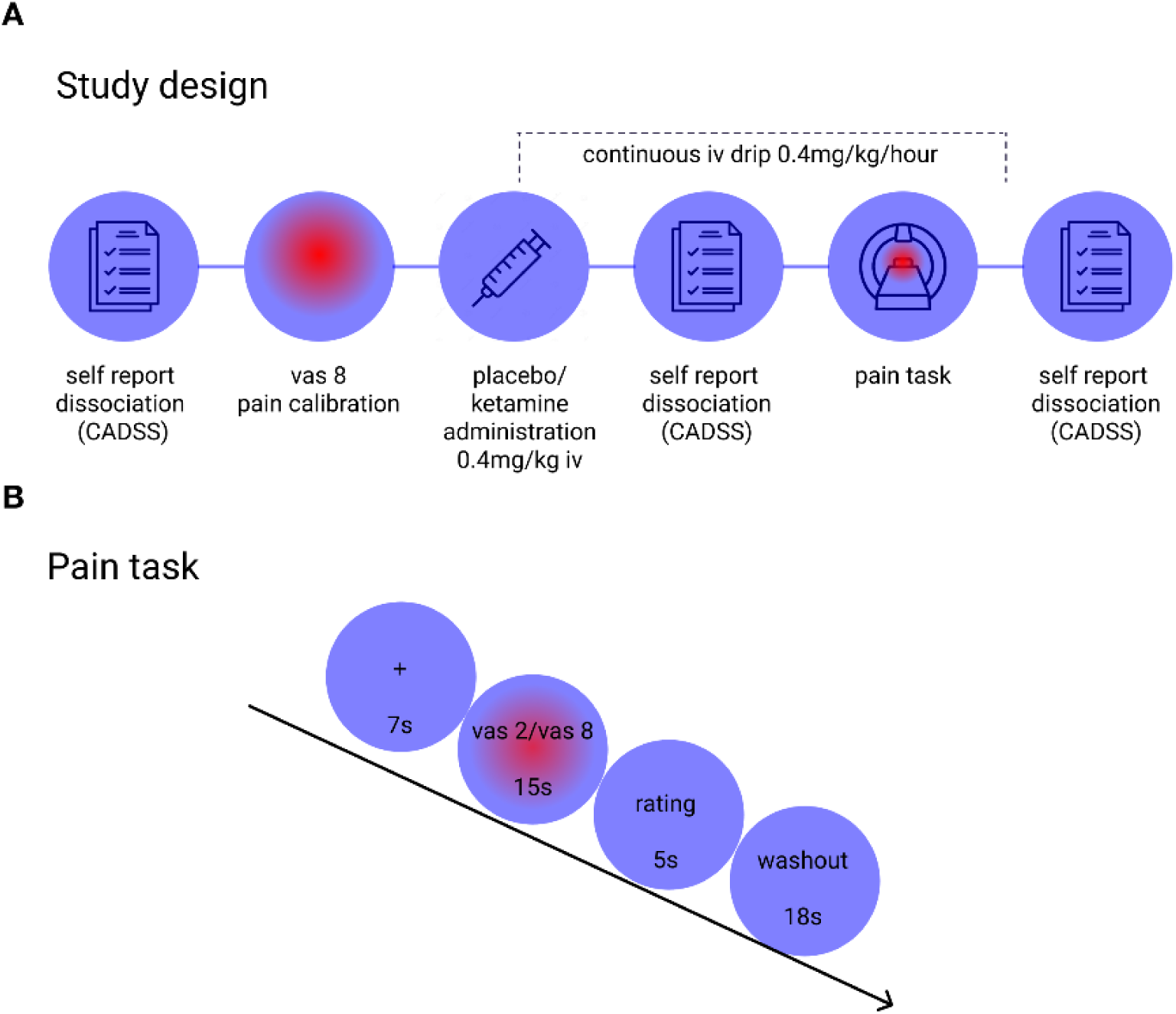
Experimental procedure and fMRI task. (A) Overview of the experimental procedure. (B) Experimental paradigm: Each trial of the pain task lasted 45s and started with a 7s fixation cross. Participants then experienced a sensation they previously rated as either painful (VAS 8) or non-painful (VAS 2) for 15s. Afterwards, they rated their experience on a five-point scale. To mitigate carryover effects between trials, an 18s visuospatial control task was introduced, during which participants viewed an arrow pointing either left or right and identified its direction.

We hypothesized that, compared to placebo, ketamine would reduce pain intensity and increase feelings of dissociation. Drawing on previous studies (e.g.,^25^), we anticipated that the analgesic effect of ketamine would be associated with decreased activity in multiple brain regions known to mediate pain perception, including somatosensory areas, the anterior insula, and the dACC. We then examined whether changes in neural activation related to ketamine’s analgesic effects also correspond to individual differences in dissociation ratings. Furthermore, we examined whether dissociation corresponds with changes in brain network connectivity, focusing on potential disintegration within the DMN and SN. To this end, we characterized the brain network connectivity affected by ketamine and investigated whether these changes correlated with individual differences in dissociation.

## Results

### Behavioral results

#### Effect of ketamine on pain perception

The pain task^31^ involved “pain” and “no pain” conditions, delivered as thermal stimulation to the right shin (L4 dermatome). Participants rated pain intensity on a ten-point scale after each trial. At the beginning of each session, prior to drug administration, painful and non-painful temperatures were individually calibrated to maintain consistent stimulus intensity^32–34^; see supplementary material for details). Out of 33 participants who completed the full study, 5 participants did not complete the pain task in both sessions, either due to technical difficulties or because they requested to stop. Across sessions, participants’ calibration results were comparable (non-painful temp; placebo = 41.7 ± 1.94°C, ketamine = 41.8 ± 1.98°C, t(31) = -.12, p = .91; painful temp; placebo = 46.1 ± 1.52°C, ketamine = 45.9 ± 1.72°C, t(31) = 1.16, p = .25). Inside the scanner, trials lasted 45 seconds, starting with a 7-second fixation cross, followed by 15 seconds of thermal stimulation (1.5 seconds for temperature ramping up, 12 seconds at target temperature, 1.5 seconds for temperature ramping down), during which participants focused on the sensation (Figure 1.B).

To investigate ketamine’s impact on pain perception, we employed a mixed-effects model with session (ketamine/placebo) and stimulus intensity (painful/non-painful) as predictors. As anticipated, the painful stimulus elicited a pronounced subjective perception of pain compared to the non painful stimulus (F(1,27) = 239.74, p < .001), and the administration of ketamine alleviated pain perception, as indicated by the main effect of session (F(1,54) = 34.88, p < .001) and the session X intensity interaction (F(1,54) = 11.22, p < .001). Post hoc analysis indicated that ketamine specifically influenced pain ratings induced by the painful stimulus (t(54) = 6.55, p < .001) but had no significant impact on the non painful stimulus (t(54) = 1.81, p = .076) (Figure 2.A).

**Figure 2.**
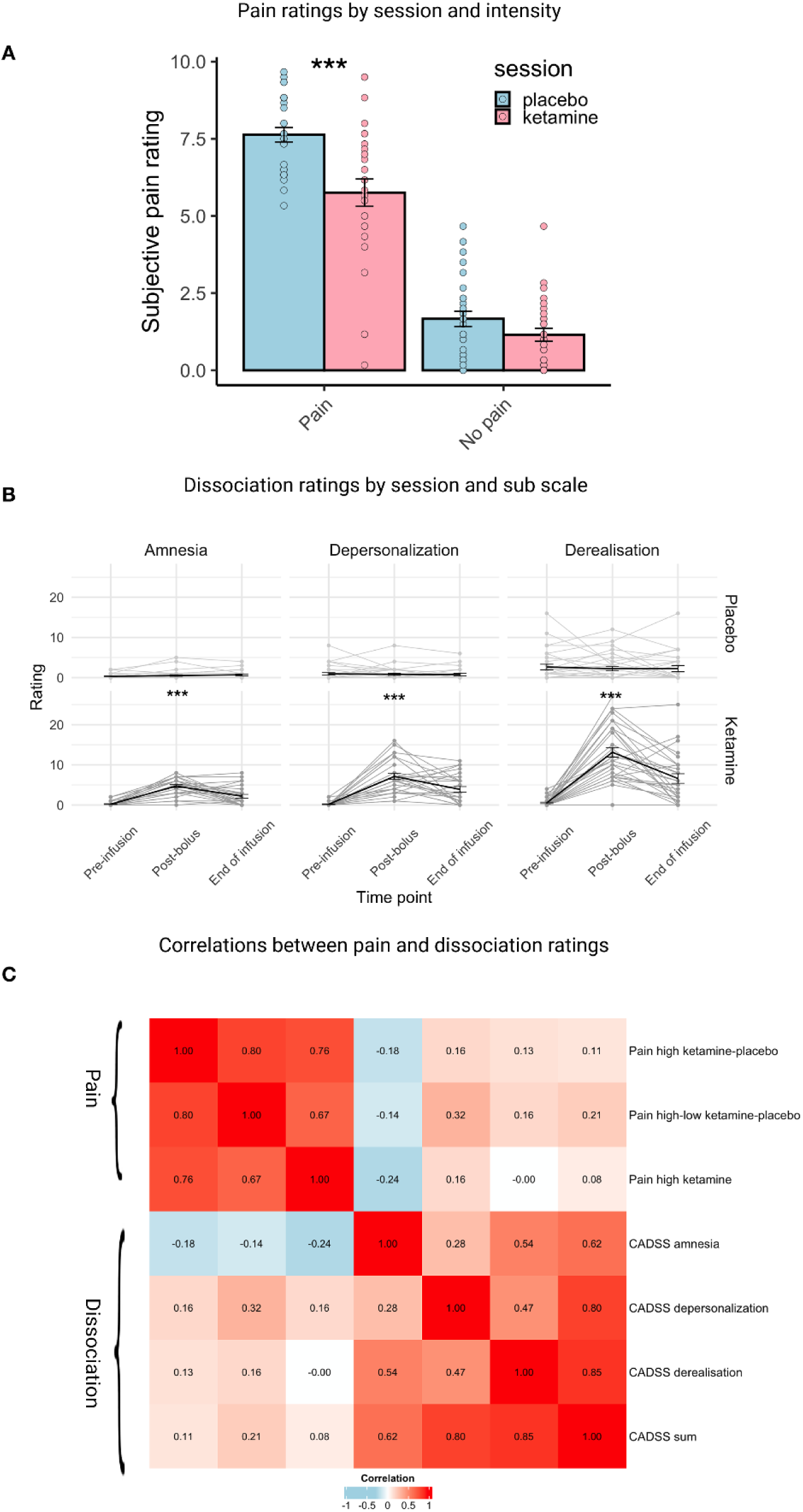
Behavioral results. (A). Subjective pain ratings for painful and non-painful stimuli during the pain task. The ketamine intervention led to a reduction in subjective pain intensity for the painful condition, but not for the non-painful condition. Points represent individual mean ratings for each subject in each condition. Bars represent the group means, and error bars indicate Standard Error of the Mean (SEM). (B). Dissociation ratings: Dissociation levels were assessed using the CADSS at three intervals during each session: pre infusion—60 minutes before the infusion began, ‘post bolus’—10 minutes after the infusion started, and ‘end of infusion’—30 minutes following the end of the infusion. Each set of three points connected by lines represents the subjective dissociation ratings across these time points for a given subscale of the CADSS questionnaire (amnesia, depersonalization, derealization). Thicker black lines indicate the mean ratings for each subscale. Error bars represent the SEM. (C). Spearman correlation coefficient matrix: This matrix displays the correlation coefficients between self-reported measures of pain and dissociation. Darker shades of red indicate stronger positive correlations, and darker shades of blue signify stronger negative correlations. * p < .05, **p < .005, *** p < .001

#### Effect of ketamine on dissociative state

Dissociation was measured using CADSS at 3 points at every session: ‘pre infusion’ – 60 minutes before infusion started, ‘post bolus’ – 10 minutes after infusion started, and ‘end of infusion’ – 30 minutes after infusion was terminated. To test the effect of ketamine on subjective feelings of dissociation we used a linear mixed-effects model that included session, time point, CADSS subscale (depersonalization, derealization, amnesia), and their interactions as predictors. Ketamine induced a strong subjective feeling of dissociation across subscales indicated by a main effect for session (F(1,32.07) = 57.44, p < .001). Peak effect was evident 10 minutes after infusion started indicated by session X time point interaction (F(2,477.73) = 109.51, p < .001). Ketamine impacted all dissociation subscales, with a significant difference observed between the ‘pre infusion’ and ‘post bolus’ time points in the ketamine session (all p’s < .001), while no such difference was found in the placebo session (all p’s > .9). (Figure 2.B). See Supplemental tables S2, S3 and S4 for the full output of this model.

#### Correlation between pain and dissociation ratings

To explore our primary hypothesis regarding a potential relationship between dissociation and pain perception during the ketamine session, we conducted a series of Spearman correlation analyses. Dissociation was assessed at the post-bolus peak timepoint, using scores from the three CADSS subscales and the total CADSS score. These were correlated with three pain-related indices: the change in pain ratings between ketamine and placebo sessions, the difference between pain and no-pain ratings, and subjective pain ratings for high-intensity stimuli during the ketamine session. Dissociation ratings were generally intercorrelated (ρ = .47–.85, all p’s < .011), except for amnesia and depersonalization, which were only weakly associated (ρ = .28, p = .151). Similarly, pain indices were significantly correlated (ρ = .67–.80, all p’s < .001), indicating internal consistency within each domain. We found no significant correlations between dissociation and pain ratings (ρ = –.24–.32, all p’s > .09), suggesting no clear association between the subjective experience of dissociation and pain perception in the context of ketamine administration (Figure 2.C).

### fMRI results

#### Pain response: univariate analysis

##### Placebo session

A whole-brain analysis contrasting painful vs. non-painful task conditions during the placebo session (n=32) revealed enhanced activation in brain regions previously associated with pain processing. These regions include the anterior insula, dACC, and dorsolateral prefrontal cortex (dlPFC), as well as S1, and supramarginal gyrus/secondary somatosensory cortex (S2; Figure 3.A and Table 1). The findings from this analysis were significant at a voxel-level threshold of p<.001, corrected for false discovery rate (FDR) cluster-level threshold of .05.

**Figure 3.**
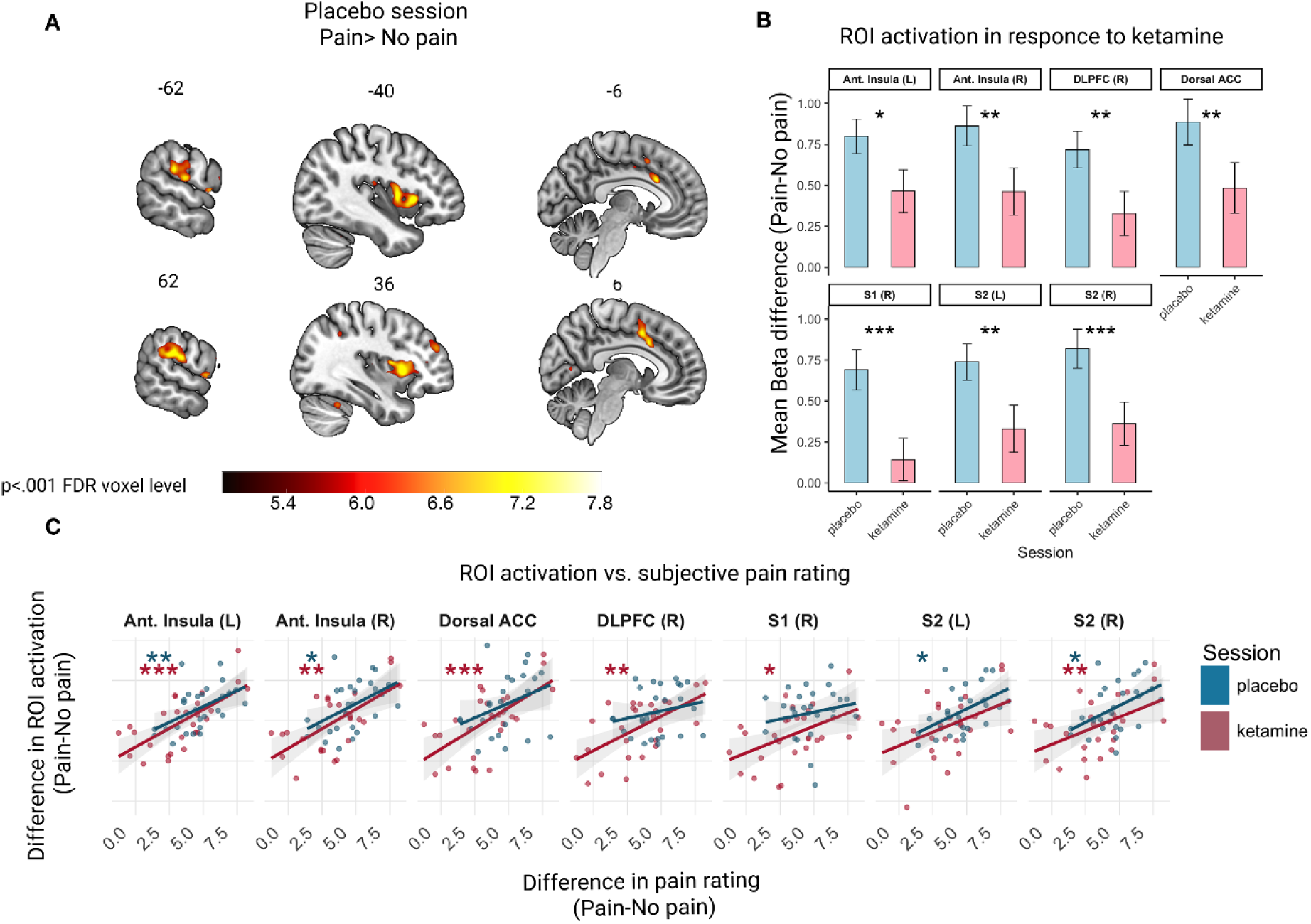
BOLD signal activation during the pain task. (A) Activation map for the contrast between painful and non-painful stimuli in the placebo session. Yellow indicates higher t-values, indicating that voxels exhibited greater activation during the painful condition compared to the non-painful condition. (B) Regions of interest (ROIs) analyzed for differences in activation patterns between the placebo and ketamine sessions were selected based on the statistical map from panel A. Panels display mean beta values for each ROI with painful-non painful task conditions in both placebo and ketamine sessions. Error bars represent the SEM. (C). Relationship between differences in subjective pain ratings and ROI activation (painful – non-painful trials) across sessions. Scatter plots show individual participants’ data points (placebo in blue, ketamine in pink) for each ROI. The x-axis represents the difference in subjective pain ratings between high and low pain conditions, and the y-axis represents the corresponding difference in ROI activation. Shaded bands indicate 95% confidence intervals for linear regressions. Asterisks denote significance of Spearman correlations. Correlation significance is color-coded by session. * p < .05, **p < .005, *** p < .001

**Table 1.** Pain-evoked brain activation during the placebo session. All clusters shown survived a voxel-level threshold of p < .001 FDR-corrected and a cluster-level threshold of p < .05, FDR-corrected.

| Brain region | L/R | Cluster size | Peak p-value | MNI coordinates |  |  |
| --- | --- | --- | --- | --- | --- | --- |
|  |  |  |  | x | y | z |
| Anterior insula /Opercular Cortex | R | 1090 | <.001 | 56 | 8 | 4 |
| Anterior insula / Opercular Cortex | L | 709 | < .001 | -44 | 2 | 6 |
| S2 | R | 329 | < .001 | 62 | -32 | 26 |
| ACC |  | 313 | < .001 | 4 | 8 | 46 |
| S2 | L | 229 | < .001 | -56 | -30 | 24 |
| Dorsolateral Prefrontal cortex | R | 68 | < .001 | 36 | 48 | 26 |
| Cerebellum | L | 23 | <. 001 | 32 | -54 | -28 |
| S1 | R | 16 | < .001 | 52 | 6 | 44 |
| Dorsal Striatum | L | 16 | < .001 | -26 | 6 | 0 |

##### Effects of Ketamine on univariate pain related brain activation

To examine the effects of ketamine on regional neural activations, we analyzed all significant cortical clusters extracted from the contrast between painful and non-painful conditions during the placebo session. A linear mixed-effects model was utilized, incorporating task condition (painful vs. non-painful), session (placebo vs. ketamine), and ROI as predictors. The model’s dependent variable was the mean beta values for each participant in each session for each ROI (n=28).

We observed a significant main effect of intensity (F(1,702) = 124.27, p < .001), indicating stronger brain activation in the painful compared to the non-painful condition. There was also a significant effect of ROI (F(6,702) = 8.11, p < .001), suggesting that different brain regions were activated to varying degrees. A main effect of session (F(1,702) = 43.05, p < .001) indicated that ketamine reduced overall activation magnitude. Additionally, we observed a significant interaction between intensity and session (F(1,702) = 16.45, p < .001) potentially indicating that ketamine had a differential effect on brain activation as a function of stimulus intensity. Post hoc analyses revealed that the difference in activation strength between pain and no-pain conditions was reduced in the ketamine session compared to placebo. This effect was observed across all pain-sensitive ROIs, including the left anterior insula (t(338) = 2.46, p = .014), right anterior insula (t(338) = 2.95, p = .003), dACC (t(338) = 2.97, p = .003), right dlPFC (t(338) = 2.87, p = .004), right S1 (t(338) = 4, p < .001,) left secondary somatosensory cortex (S2 L) (t(338) = 3, p = .003), and right secondary somatosensory (S2 R) (t(338) = 3.36, p < .001) (Figure. 3.B and Supplemental Tables S5, S6).

##### Correspondence between univariate pain-related brain activation and pain ratings

To assess whether activation in pain-related ROIs corresponded to participants’ subjective experience of pain, we computed within-session Spearman correlations between differences in pain ratings (pain – no pain) and corresponding differences in ROI activation (pain – no pain), separately for the placebo and ketamine sessions. In the placebo session, activation in pain-responsive ROIs was positively correlated with reported pain intensity (mean ρ = .369, range: .134–.528). Significant correlations were observed in four ROIs: left anterior insula (ρ = .528, p = .005), right anterior insula (ρ = .463, p = .015), left S2 (ρ = .460, p = .016), and right S2 (ρ = .427, p = .026). Correlations in other ROIs, including the dACC, right dlPFC, and right S1, were positive but did not reach significance. In the ketamine session, the overall strength of correlations was numerically higher (mean ρ = .532, range: .344–.656), and six of seven ROIs showed significant associations with pain ratings. These included the left anterior insula (ρ = .656, p < .001), right anterior insula (ρ = .561, p = .002), dACC (ρ = .651, p < .001), right dlPFC (ρ = .570, p = .002), right S1 (ρ = .448, p = .019), and right S2 (ρ = .491, p = .009). The left S2 showed a nonsignificant but positive correlation (ρ = .344, p = .079). These results indicate that subjective pain intensity is univariately represented in most pain-related ROIs, and this relationship is consistent across both placebo and ketamine administration (see Figure 3.C and Supplemental Table S7).

##### Correspondence between univariate pain-related brain activation and dissociation ratings

To examine whether pain-related brain activation under ketamine corresponded specifically to subjective pain experience, or whether it might also reflect dissociative symptoms, we tested whether dissociation ratings explained additional variance in activation of pain-sensitive ROIs. For each ROI, we computed the difference in activation between high- and low-pain conditions in the ketamine session. We first modeled this activation difference as a function of subjective pain ratings (pain-no pain in the ketamine session) and then assessed whether including dissociation ratings, calculated as the change from placebo to ketamine in CADSS subscales (Depersonalization, Derealization, Amnesia), significantly improved model fit. Pain ratings significantly predicted activation in most pain-sensitive regions (all p’s < .05), with the exception of right S1 (p = .09). However, dissociation ratings did not account for additional variance in any ROI (all p > .63; see Supplemental Table S8). These findings suggest that ketamine’s analgesic effects are reflected in pain-related brain activation independently of its dissociative effects.

##### Pain response: multivariate analysis

To further explore the relation between reported subjective analgesia exerted by ketamine and objective brain measures, we examined ketamine’s impact on the expression of a multi-voxel pattern previously validated as a biomarker for the subjective intensity of pain: the Neurologic Pain Signature (NPS)^35^. Unlike approaches that treat voxel-level brain activity as the outcome, this multi-voxel biomarker uses subjective pain intensity as the variable to be predicted or explained by distributed patterns of brain activity. Once validated, such markers can serve as objective indicators of the subjective experiences they quantify^36^. A linear mixed-effects model including task condition (painful vs. non-painful) and session (placebo vs. ketamine) revealed a significant main effect of condition (F(1,26) = 42.91, p < .001), indicating that NPS responses were higher during painful compared to non-painful stimuli across both sessions. There was also a significant main effect of session (F(1,26) = 4.71, p = .039), as well as a condition-by-session interaction (F(1,26) = 5.43, p = .028), suggesting that ketamine modulated NPS responses, particularly in the painful condition. NPS responses to the painful condition were significantly lower under ketamine than placebo (t(41.4) = 3.04, p = .004), with no difference in the non-painful condition (t(41.4) = 0.73, p = .47). Nonetheless, the NPS maintained its specificity and could distinguish between painful and non-painful both in the placebo (t(49) = 6.6, p < .001) and ketamine sessions (t(49) = 3.74, p < .001) (see Figure. 4.B).

**Figure 4.**
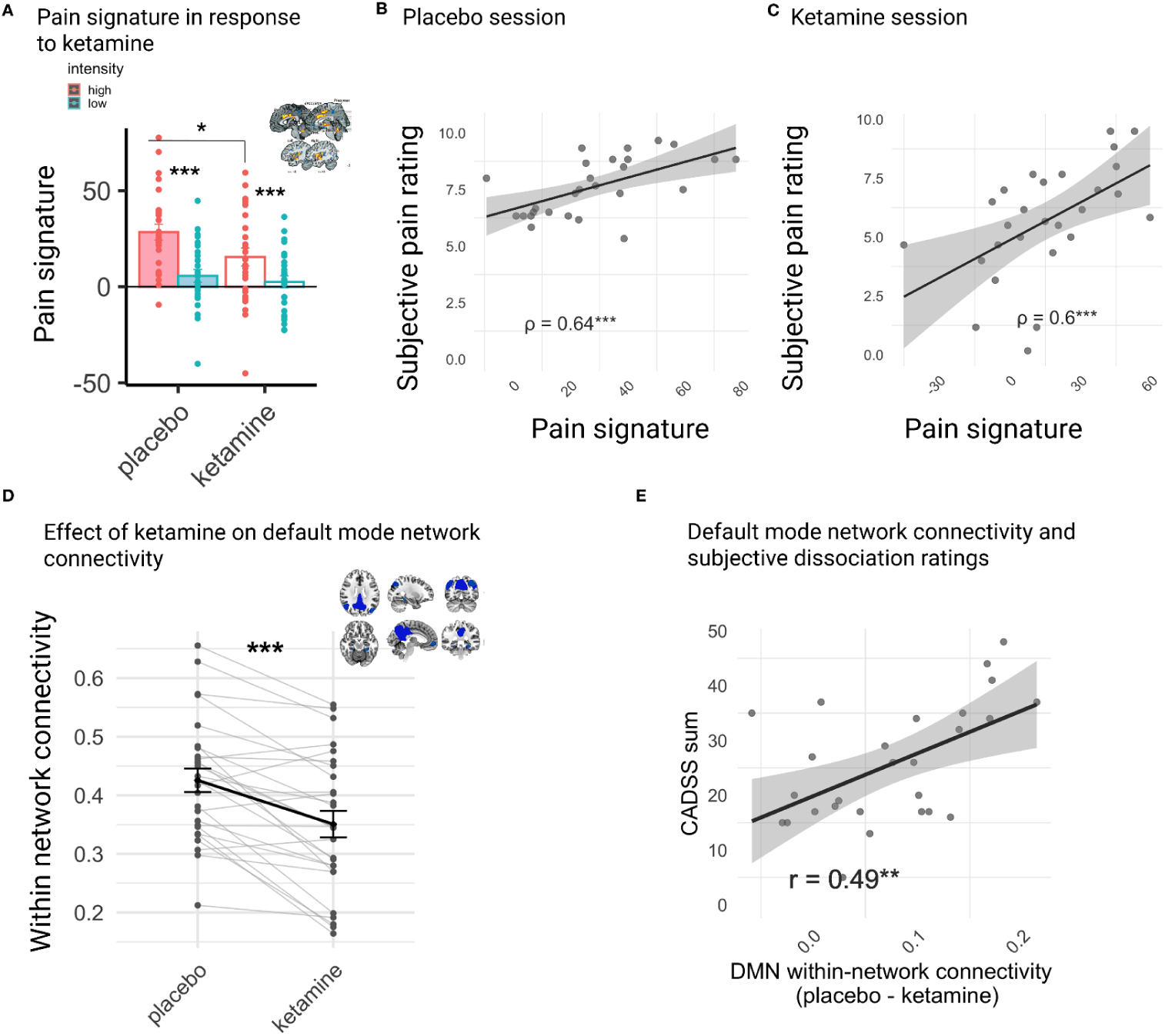
Neural correlates of ketamine’s analgesic and dissociative effects. (A) The Neurologic Pain Signature (NPS^35^) biomarker predicted a stronger pain response in the painful condition during the placebo session compared to the ketamine session. In both sessions, this biomarker predicted a stronger pain response in the painful condition compared to the non-painful condition. Points represent individual mean NPS responses for each subject in each condition. Bars represent group means, and error bars indicate the SEM. (B,C) During both placebo and ketamine sessions, there was a significant correlation between subjective pain ratings and NPS response during the painful condition. Shaded areas depict 95% confidence intervals. (D) Ketamine led to a reduction in within-network connectivity of the Default Mode Network (DMN). Each pair of points connected by lines represents the mean person correlation between all brain areas constituting the DMN (presented in blue at the top right of panel D) for each session. Thicker black lines indicate the mean correlation at the group level. Error bars represent SEM. (E) Subjective dissociation ratings were correlated with a reduction in connectivity within DMN nodes. Shaded areas depict 95% confidence intervals.

##### Correspondence between multivariate pain-related brain activation and pain ratings

We then examined whether the established relationship between subjective pain ratings and NPS responses was preserved following ketamine administration. To this end, we computed Spearman’s correlations between pain ratings and NPS expression values within each session. Results showed robust correlations during the painful condition in both the placebo (ρ = .64, p < .001) and ketamine (ρ = .60, p < .001) sessions, indicating that the NPS remained a reliable marker of subjective pain even under ketamine (Figure 4.B). Importantly, adding dissociation ratings to the model did not account for additional variance in NPS expression beyond that explained by pain ratings alone (p = .28; see Supplemental Table S9), suggesting that multivariate pain-related brain activity under ketamine is not influenced by dissociative experience.

##### Neural correlates of dissociative states

Both univariate and multivariate analyses of pain-related brain activation under ketamine did not correlate with subjective dissociation ratings. Nevertheless, we were interested in examining if changes in neural indices previously linked to ketamine-induced dissociation could explain individual variability in our sample and, if so, whether these indices would be uniquely linked to dissociation ratings or would also correspond to ketamine’s analgesic effect. Most studies linking brain signals to dissociation have focused on functional connectivity, particularly in the DMN and SN networks^26–30^. Building on prior evidence that ketamine modulates these systems, we examined functional connectivity across five brain networks during the pain task: the DMN, SN, Frontoparietal Network (FPN), Sensorimotor Network (SMN), and Dorsal Attention Network (DAN) (see Methods for further details).

Network-level connectivity was assessed using a built-in function in the CONN toolbox, which calculates within-network connectivity by averaging Fisher-transformed correlation coefficients between all ROI pairs within each network. The most pronounced ketamine-related effect was observed in the DMN (t = -5.38, p < .001). The impact on other networks was not statistically significant (all p’s > .343; see Supplementary Figure S.1). Then, to evaluate whether ketamine-induced changes in DMN connectivity correspond to individual differences in dissociation ratings, we calculated the differences between ketamine and placebo DMN within-network connectivity values for each participant and calculated the Spearman correlation between these differences and peak dissociation ratings post-bolus. We observed a significant correlation between diminished within-network DMN connectivity and higher dissociation ratings (CADSS sum: ρ = .489, p<.01; see Figure 4.E). When examining correlations with CADSS subscales, significant correlations were observed with derealization (ρ = .58, p = .005) and amnesia (ρ = .374, p = .05) subscales, whereas no significant correlation was found with the depersonalization subscale (ρ = .235, p = .228). Additionally, we found no significant correlation between changes in within-network DMN connectivity and reduction in pain ratings between sessions (ρ = -.055, p = .783), nor with high pain ratings during the ketamine session (ρ = -.072, p = .718). This indicates that the disintegration of the DMN following ketamine intervention is linked to subjective feelings of dissociation, but not to ketamine-induced analgesia.

## Discussion

This study addressed whether the dissociative properties of ketamine explain its ability to ameliorate pain. This question reflects a broader debate about whether the subjective experiences elicited by psychoactive drugs are necessary for therapeutic outcomes^6,7^. Following ketamine administration, participants reported reduced pain intensity, which was accompanied by decreased activation in a pain-responsive network of brain regions including the insula, dACC, and secondary somatosensory cortex. Concurrently, we observed a strong dissociative experience linked to the disintegration of the DMN. Importantly, we observed no correlation between the intensity of dissociative experiences and the analgesic effects of ketamine at both the behavioral and neural levels. These results inform an ongoing debate, which has produced mixed results thus far regarding the relationship between ketamine’s analgesic and dissociative effects. Our results suggest that these two phenomena may operate independently of each other.

Pain is a multifaceted, subjective experience that includes sensory, affective, attentional, and cognitive components^17,37,38^. Correspondingly, the experience of pain is represented in the brain by a network of regions that encode its various dimensions. For example, somatosensory regions code for properties such as stimulus location and duration^39–41^ while regions such as the anterior insula and dACC may code for pain-related expectations, motivations and subjective aversion^42–45^. Whether ketamine exerts its analgesic effect through modulation of representations related to the sensory properties of the nociceptive stimulus, or by affecting the modulation of evaluation or attentional processes is still an open question.

In this study, ketamine induced a uniform reduction in pain-related brain activation across regions that demonstrated positive pain-related activation during the placebo session, along with a decrease in the magnitude of the multi-voxel NPS response. This observation aligns with previous studies on ketamine^25,46^, which reported decreased pain-related activation across brain regions encoding both the sensory and affective dimensions of pain. Importantly, despite these reductions, ketamine did not alter the correlation between brain activation and the subjective experience of pain. Correlation with self-reported pain was observed in the activation of brain areas such as the anterior insula and dACC, as well as in the multivoxel expression of the NPS, in both the ketamine and placebo sessions. This suggests that while ketamine may decrease the overall magnitude of the pain response, it does not inherently change the neural representation of pain. The consistent correlation between brain activation and subjective pain ratings across both sessions indicates that the processes of evaluating or monitoring pain intensity remain stable under the influence of ketamine. These results, cannot arbitrate between top-down and bottom-up analgesia. However, one mechanism that might be consistent with the data presented here is that ketamine reduces the overall nociceptive input to the cortex, potentially through spinal mechanisms^47^. This idea aligns with findings from animal studies, which show that ketamine disrupts nociceptive transmission between the spine and the supraspinal pain systems^48^. At the molecular level, the modulation of this process might be mediated by the NMDA receptor, which is a key antagonistic target of ketamine^49^. NMDA receptor activation can lead to the amplification of nociceptive signals at the spinal level, resulting in pain hypersensitivity^50–53^. This increased sensitivity can manifest through modulation of the temporal summation of pain, where a stimulus delivered at constant intensity evokes increased pain reactions. Accordingly, effects on temporal summation could be reversed by using NMDA antagonists^54^. Notably, however, NMDA may not be the only signaling pathway through which ketamine can reduce nociceptive signaling. The increase in serotonergic signaling induced by ketamine might also inhibit incoming nociceptive signals to the cortex^55,56^.

The DMN has long been the focus of studies examining the effects of mind-altering drugs on changes in the sense of self^57–59^. The DMN is predominantly active during mental simulation processes and is thought to contribute to high-level, modality-invariant representations of semantic concepts, which are key to the feeling of coherent identity and sense of self^60,61^. Accordingly, observations that psychoactive substances such as ketamine, LSD, or psilocybin create alterations in self-experience^62–68^ and lead to alterations in DMN connectivity^27,69–74^ have led to propositions that the two phenomena might be mechanistically connected^59,75–77^. Notably, the DMN has also been suggested to play a role in pain perception: it is deactivated during sensory aversive perception, and increased DMN activity is involved when shifting attention away from the pain^78^. Moreover, changes in self-related experiences have been associated with alterations in pain perception. For instance, mindfulness meditation has been linked to decreased pain catastrophizing^79,80^, less negative appraisal of pain^81^, and reduced unpleasantness of pain^79^. Notably, however, a recent meta-analysis has demonstrated that the insula, ACC, PFC, and amygdala are the brain regions most consistently associated with the effects of meditation on pain responses^82^. This may indicate that even with manipulations targeting self-related functioning and valuation processes, the neural regions involved in pain relief are mostly sections of what one may call the pain matrix, and to a lesser extent, the DMN. These findings are consistent with our data but leave an open question regarding the mechanism by which reduced pain-related brain activity is achieved.

Several recent studies have tested whether dissociation induced by ketamine is mechanistically linked to its analgesic effects—a question we also address. Two out of three studies concluded that these processes are independent^23,24^. In one study^23^, researchers used an initial ketamine infusion followed by midazolam administration to reduce dissociative effects. Both pain reduction and dissociation were predicted based on the time since the beginning of the infusion. However, adding dissociation ratings did not improve the model’s ability to predict pain reduction, and vice versa. In a follow-up study^24^, the same group replicated this result in a study design where ketamine was combined with Sevoflurane, a drug commonly used for general anesthesia. Together, these two studies support the notion that the analgesic and dissociative properties of ketamine are separable. However, in a third study^22^, researchers assessed pain thresholds and perceptual disturbances (psychedelic effects) during ketamine infusion, measuring plasma concentrations of ketamine and norketamine. They aimed to identify the drug concentration at which antinociceptive or psychedelic effects appeared. The models for ketamine’s antinociceptive and psychedelic effects were not statistically distinguishable. This led the researchers to conclude that the analgesic and psychoactive effects of ketamine are interdependent^22^. Notably, none of these studies employed brain imaging, which provides an additional level of analysis and can help disentangle the overlapping temporal aspects of ketamine’s analgesic and dissociative effects.

The finding that the two effects are independent invites broader consideration. There is ongoing debate about whether the acute mind-altering properties of psychoactive substances are essential for their clinical benefits. Efforts are underway to develop non-psychoactive analogs of mind-altering compounds that exhibit clinical benefits^83–91^, yet other studies argue that the mind-altering properties are central to these benefits. A key study by Lii et al.^92^ examined the effects of a single intravenous dose of ketamine, administered during surgical anesthesia, on the acute reduction of depressive symptoms in individuals with major depressive disorder. The study found that under such conditions ketamine was not superior to placebo. This result suggests two possible interpretations: first, the subjective psychoactive effects of ketamine may be crucial for its clinical antidepressant properties (despite the lack of a systematic correlation between its dissociative and antidepressant effects^93^). Alternatively, the psychoactive effects of the drug could influence clinical trial outcomes by affecting masking and interfering with proper blinding.

The challenges associated with incomplete masking are also relevant in the context of our study. To address potential unmasking following ketamine administration, we adopted a fixed order design, where the placebo session always preceded the ketamine session. This design has some disadvantages: Despite our efforts to minimize habituation across sessions through a same-day pain calibration procedure, participants’ growing familiarity with the pain paradigm and laboratory setting during their second visit may have influenced their neural or behavioral responses. Notably, alternative designs also pose their own challenges. For instance, in a randomized order design, participants receiving ketamine first would be aware that their next session would involve a placebo, potentially biasing their expectations and responses. Conversely, those starting with placebo might not ascertain their subsequent treatment, leading to an imbalance in blinding that could introduce further confounds. Employing an active drug as a control complicates the interpretation of brain imaging results by introducing additional variables. These dilemmas underscore the inherent difficulties in ensuring proper blinding in experiments with psychoactive drugs, a topic that has been explored in depth elsewhere^94^. An additional limitation of our study is its correlational nature. To address this, future research could consider a combination of neuroimaging and an additional pharmacological manipulation that blocks ketamine’s effects on specific receptors, such as the opioid or dopaminergic systems. In this regard, it would be particularly interesting to compare ketamine with lamotrigine, an anticonvulsant that decreases presynaptic glutamate release and has been shown to reduce ketamine-induced psychological disturbances, including dissociative symptoms, as well as attenuate ketamine’s effects on the central nervous system^95–98^. This approach could help disentangle the contributions of different ketamine signaling pathways to its complex effects on human consciousness.

To conclude, this study suggests that the analgesic and dissociative effects of ketamine are distinct and are mediated by different neural systems. This finding supports the idea that in at least some of its clinical uses, a drug’s subjective effects on consciousness can be separate from its clinical benefits. While the dissociative effects of ketamine could potentially have their own clinical advantages, they do not directly account for its analgesic properties.

## Methods

### Participants

A total of 37 participants were enrolled in the study (mean age 29.68 ± 3.21 years, 21 females). Of these, four participants completed only a single session, leaving 33 participants who underwent the two-session procedure (mean age 30.1 ± 3.09 years, 18 females). Four participants encountered technical difficulties or requested to discontinue during the fMRI pain paradigm, resulting in the exclusion of their data from certain analyses. Nevertheless, their data were incorporated into the behavioral analysis. One participant had abnormal neurological findings that were discovered during the second scan and was excluded from the analysis. All participants had a minimum of 12 years of education, no reported psychiatric or neurological disorders (including ADHD), no current use of psychoactive drugs, and were native Hebrew speakers. With the exception of one female participant, all participants were right-handed. Informed written consent was obtained from all participants in accordance with the Tel-Aviv Sourasky Medical Center institutional review board (IRB) committee guidelines (ClinicalTrials.gov Identifier: NCT02037503). Participants received monetary compensation at a rate of 50 NIS per hour. Participants were recruited through social media advertisements, and the screening process aimed to ensure the inclusion of physically and mentally healthy individuals who met the MRI scanning criteria. Individuals with psychotic disorders (including first-degree relatives with such diagnoses), cardiovascular disorders, frequent alcohol use (> once a week), cannabis use (> once a month), or other drug use (> once every 3 months) were excluded.

### Overview of the experimental design

This study was a single-blind, placebo-controlled, within-subjects experiment conducted at the Sagol Brain Institute, Sourasky Medical Center, Tel-Aviv, Israel. Upon their arrival at the laboratory, participants were presented with a comprehensive overview of the experimental protocol and provided written informed consent. Following this, they underwent a training session to acquaint themselves with the experimental tasks that would subsequently be performed within the MRI scanner. A battery of self-report questionnaires, which are beyond the scope of the current study, was then administered. Subsequently, a pain calibration procedure was conducted to establish individual pain thresholds, details of which can be found in the supplementary material. Once calibration was completed, participants were escorted to a comfortable room where qualified physicians inserted intravenous catheters. Participants then received infusions of either a placebo or ketamine in two stages: an initial bolus loading dose followed by a continuous steady drip. Further details are provided in the subsequent sections. The placebo session was consistently introduced first, with participants unaware of this sequence. Approximately ten minutes after the conclusion of the initial bolus infusion, participants responded to a verbally administered CADSS^9^, facilitated by the researcher. This was followed by the completion of a cognitive task that lies outside the scope of this paper. Following this task, participants underwent a 60-minute MRI scan comprising an anatomical scan, a double-echo gradient-echo field map sequence, and functional MRI sequences for the pain task^31,99^. As well as other fMRI tasks. Here, we report only analysis of the physical pain task, while the social pain task will be reported elsewhere. At the conclusion of the MRI scan, participants were disconnected from the IV infusion, and they subsequently addressed another set of self-report questionnaires.

### Ketamine/ placebo administration protocol

We combined a previously established ketamine induction protocol from D’souza et al.^100^ with insights gathered from pilot sessions conducted in our lab. Our goal was to induce a noticeable psychoactive effect while ensuring that participants remained capable of performing behavioral tasks, comprehending instructions, and maintaining full awareness and communicative abilities. To achieve this, we administered an initial bolus of .4 mg/kg IV over 10 minutes, followed by a continuous drip of .4 mg/kg/h using an infusion pump. This approach provided a sustained and stable psychoactive experience. For the placebo condition, participants received an IV infusion of saline (.9% NaCl) in a volume equivalent to that of the ketamine solution.

### Self-reported dissociation

To assess dissociation, we used the CADSS, a well-established tool in this context^9,101^. The version of CADSS that was used comprises 23 items, each rated on a scale from 0 (no effect) to 4 (extreme effect). These items evaluate various aspects, including changes in body sensation, perception of time and environment, memory function, and alterations in the sense of reality. The CADSS organizes these items into three subscales: depersonalization, derealization, and amnesia. Numerous randomized-controlled trials have utilized CADSS to evaluate the acute psychoactive impacts of ketamine^102–105^.

### Physical pain fMRI task

To investigate physical pain, we employed a previously established protocol^31^, which also incorporated an additional condition assessing mental pain, though the latter is beyond the scope of this paper. The task featured two conditions: pain and no pain, each delivered via thermal stimulation applied to the participant’s right leg, specifically the lateral aspect of the shin area (L4 dermatome). Following each trial, participants rated pain intensity on a ten-point scale, with higher scores indicating more intense pain. To ensure consistency in subjective stimulus intensity across participants, we individually calibrated painful and non-painful temperatures based on each participant’s reaction, separately for each session, a method previously employed^32–34^, see supplementary material for additional detail). Each trial lasted 45-s and began with a 7-s fixation cross. Subsequently, participants experienced a 15-s thermal stimulation period. During this period, participants viewed a fixation cross and focused on the sensations they experienced as a hot (pain) or warm (no pain) stimulus was applied (1.5-s temperature ramp up/down, 12 s at peak temperature). They then rated the pain they experienced using a ten-point scale (0 = not painful; 10 = very painful). To reduce carryover effects between trials, participants then performed an 18-s visuospatial control task in which they saw an arrow pointing left or right and were asked to indicate which direction the arrow was pointing.

### Visuospatial control task

As an integral component of the pain task, each trial included an 18-s visuospatial control task. During this task, participants observed an arrow pointing either left or right and indicated the arrow’s direction. This task served two purposes: (a) mitigating carryover effects from the painful stimulus and (b) ensuring participant engagement, responsiveness, and verifying similar simple motor performance between the ketamine and placebo sessions.

### fMRI data acquisition

Scans were conducted using a Siemens 3T Prisma Magnetom VD13 echo speed scanner equipped with a 20-channel head coil at the Wohl Institute for Advanced Imaging, Tel-Aviv Sourasky Medical Center. Structural scans included a T1-weighted 3D axial spoiled gradient echo (SPGR) pulse sequence (TR/TE = 1,860/2.74 ms, flip angle = 8°, voxel size = 1 x 1 x 1 mm, field of view = 256 × 256 mm, slice thickness = 1 mm). Functional whole-brain scans for all fMRI tasks followed an interleaved bottom-to-top order and utilized a T2*-weighted gradient echo-planar imaging pulse sequence (TR/TE = 2500/30 ms, flip angle = 82°, voxel size = 2.0 x 2.0 x 3.0 mm, field of view = 220 × 220 mm, slice thickness = 3 mm, 42 slices per volume). 444 were acquired. Multi-echo GE images were obtained for field mapping before the functional scan, with geometrical and orientational correspondence to the functional runs. Additional sequence parameters included TE = [4.92,7.38] ms, TR = 400 ms, FA = 60, and voxel size: 3.4×3.4×3.0 mm.

### Preprocessing and GLM analysis

Functional MRI data were preprocessed using *fMRIPrep*^106^; see Supplementary Materials for full details. Preprocessed images were spatially smoothed with a 6 mm FWHM kernel using SPM12. First-level analyses modeled boxcar regressors convolved with the canonical hemodynamic response function for pain stimulation (15 s), rating (8 s), a visuospatial control task (18 s), and fixation (7.5 s). The TR was 2.5 seconds. Nuisance regressors included six head motion parameters and their derivatives, quadratics, and squared derivatives (totaling 24), framewise displacement (FD), and 18 physiological regressors (e.g., heart rate and respiration). Volumes with FD > .4 mm were marked as motion outliers and scrubbed^107^. First-level models were estimated using SPM12, and individual contrast maps were carried forward into group analyses.

### Neurological Pain Signature (NPS)

We used previously developed multivariate spatial patterns that depict neural activations caused by physical pain. This pattern is named neurological pain signature (NPS), shown to reliably and selectively predict physical pain intensity in human participants. This pattern signature involves to a significant extent brain regions such as the dACC, anterior insula, medial thalamus, and somatosensory cortices^35^. Application of the NPS to a new dataset involves multiplying NPS-per voxel weights in each activation map in the new dataset (within-person and condition) and reducing the product to a weighted average. The procedure has been used in several past studies^108–111^

### Functional connectivity

We conducted a functional connectivity analysis via the CONN toolbox (CONN 20a)^112^. For this analysis, we first imported into CONN the data that was pre-processed in fMRIPrep based on the SPM.mat files that were constructed for the 1st-level GLM. Thus, the confound regressors were similar to those used in the neural activation GLMs, apart from the aCompCor regressors which we re-computed in CONN. The following pre-processing steps were performed with the CONN toolbox. First, masks of white matter, grey matter, and cerebrospinal fluid were produced by performing segmentation on the structural image of each subject. In addition to the cofound regressors imported from the SPM.mat, the following confounds were regressed out during the denoising step: (1) the first five principal components of the CSF and white matter signals, based on the aCompCor method, and (2) task-related BOLD signals by performing linear detrending. Bandpass temporal filtering was performed to remove slowly fluctuating signals (.0008 Hz) such as scanner drift. To account for variance explained by all experimental conditions, all task conditions were defined as experimental conditions in the setup phase in CONN.

### Network connectivity analysis

To identify relevant functional networks, we performed group-level independent component analysis (ICA) using the CONN toolbox^112^, following standard procedures^113^. Extracted components were labeled using the built-in overlap tool comparing each component with a canonical brain-network parcellation^114^ . This process identified five networks of interest: the DMN, SN, FPN, SMN, and DAN. Within-network connectivity was computed for each participant and session using the CONN toolbox’s built-in function. Specifically, we extracted BOLD time series from the pain task and calculated the pairwise Pearson correlation coefficients between all ROI pairs within each network. The Fisher-transformed average of these correlation coefficients was used as a measure of within-network connectivity. Between-session differences (ketamine vs. placebo) were assessed. Further information on ROIs, network definitions, and MNI coordinates is provided in the Supplementary Materials.

### Statistical analysis

We used R version 4.1.0 and MATLAB R2021a for statistical analyses. Mixed-effects regression models were conducted using the optimizer “bobyqa” with one million model iterations in the *afex* package version 1.3-1^115^. Models included the maximal random-effects structure (i.e., random intercepts, slopes, and their correlations across fixed effects for each subject) to minimize Type I error^116^. If a model didn’t converge, we reduced the random-effects structure. For all linear models, the significance of fixed effects was determined by an *ANOVA* using the Kenward-Roger method to calculate degrees of freedom. Post hoc comparisons were conducted using the emmeans package (version 1.8.7). Whole-brain fMRI analyses applied conservative correction thresholds: voxel-wise FDR at p < .001 and cluster-level FDR at p < .05. In post hoc ROI analyses, no correction for multiple comparisons was applied.

## Author contributions

Noam Goldway. conceived and designed the study, led the investigation, data collection, curation, and formal analysis, and drafted the original manuscript. Talma Hendler provided supervision, secured funding, and contributed to statistical oversight. Itamar Jalon contributed to data preprocessing, formal analysis and edited the manuscript. Yotam Pasternak, Roy Sar-El and Dan Mirelman were responsible for participant recruitment, supervision, and study procedure management Noam Sarna and Nili Green. contributed to data collection and management. Yara Agbaria supported statistical analysis and preprocessing. Haggai Sharon supervised project development, provided statistical consultation, and critically reviewed manuscript drafts. All authors reviewed and approved the final manuscript.

## Acknowledgments

We thank Shira Niv, Shiba Sitry, Yuval Koryto, and Rony Hirschhorn for their help with data collection for this manuscript.

## Data availability

The code and data supporting the findings of this study are publicly available via the Open Science Framework (OSF): https://osf.io/k4wy5/

## References

1. Nutt, D. & Carhart-Harris, R. The Current Status of Psychedelics in Psychiatry. JAMA Psychiatry 78, 121–122 (2021).

2. Elman, I., Pustilnik, A. & Borsook, D. Beating pain with psychedelics: Matter over mind? Neurosci. Biobehav. Rev. 134, 104482 (2022).

3. Krystal, J. H., Abdallah, C. G., Sanacora, G., Charney, D. S. & Duman, R. S. Ketamine: A Paradigm Shift for Depression Research and Treatment. Neuron 101, 774–778 (2019).

4. Bell, R. F. & Kalso, E. A. Ketamine for pain management. Pain Rep 3, e674 (2018).

5. Yaden, D. B., Earp, B. D. & Griffiths, R. R. Ethical Issues Regarding Nonsubjective Psychedelics as Standard of Care. Camb. Q. Healthc. Ethics 31, 464–471 (2022).

6. Olson, D. E. The Subjective Effects of Psychedelics May Not Be Necessary for Their Enduring Therapeutic Effects. ACS Pharmacol Transl Sci 4, 563–567 (2021).

7. Yaden, D. B. & Griffiths, R. R. The Subjective Effects of Psychedelics Are Necessary for Their Enduring Therapeutic Effects. ACS Pharmacol Transl Sci 4, 568–572 (2021).

8. Spiegel, D. et al. Dissociative disorders in DSM-5. Depress. Anxiety 28, E17–45 (2011).

9. Bremner, J. D. et al. Measurement of dissociative states with the Clinician-Administered Dissociative States Scale (CADSS). J. Trauma. Stress 11, 125–136 (1998).

10. Bohus, M. et al. Pain perception during self-reported distress and calmness in patients with borderline personality disorder and self-mutilating behavior. Psychiatry Res. 95, 251–260 (2000).

11. Ludäscher, P. et al. Pain sensitivity and neural processing during dissociative states in patients with borderline personality disorder with and without comorbid posttraumatic stress disorder: a pilot study. J. Psychiatry Neurosci. 35, 177–184 (2010).

12. Defrin, R., Schreiber, S. & Ginzburg, K. Paradoxical Pain Perception in Posttraumatic Stress Disorder: The Unique Role of Anxiety and Dissociation. J. Pain 16, 961–970 (2015).

13. Schmahl, C. et al. Pain sensitivity is reduced in borderline personality disorder, but not in posttraumatic stress disorder and bulimia nervosa. World J. Biol. Psychiatry 11, 364–371 (2010).

14. Krystal, J. H. et al. Subanesthetic effects of the noncompetitive NMDA antagonist, ketamine, in humans. Psychotomimetic, perceptual, cognitive, and neuroendocrine responses. Arch. Gen. Psychiatry 51, 199–214 (1994).

15. Hocking, G. & Cousins, M. J. Ketamine in chronic pain management: an evidence-based review. Anesth. Analg. 97, 1730–1739 (2003).

16. Persson, J. Ketamine in pain management. CNS Neurosci. Ther. 19, 396–402 (2013).

17. Bushnell, M. C., Ceko, M. & Low, L. A. Cognitive and emotional control of pain and its disruption in chronic pain. Nat. Rev. Neurosci. 14, 502–511 (2013).

18. Reddan, M. C. & Wager, T. D. Modeling Pain Using fMRI: From Regions to Biomarkers. Neurosci. Bull. 34, 208–215 (2018).

19. De Ridder, D., Adhia, D. & Vanneste, S. The anatomy of pain and suffering in the brain and its clinical implications. Neurosci. Biobehav. Rev. 130, 125–146 (2021).

20. Mashour, G. A. Ketamine Analgesia and Psychedelia: Can We Dissociate Dissociation? Anesthesiology 136, 675–677 (2022).

21. Schwenk, E. S. et al. Consensus Guidelines on the Use of Intravenous Ketamine Infusions for Acute Pain Management From the American Society of Regional Anesthesia and Pain Medicine, the American Academy of Pain Medicine, and the American Society of Anesthesiologists. Reg. Anesth. Pain Med. 43, 456–466 (2018).

22. Olofsen, E. et al. Ketamine Psychedelic and Antinociceptive Effects Are Connected. Anesthesiology 136, 792–801 (2022).

23. Gitlin, J. et al. Dissociative and Analgesic Properties of Ketamine Are Independent. Anesthesiology 133, 1021–1028 (2020).

24. Hahm, E. Y. et al. Dissociative and analgesic properties of ketamine are independent and unaltered by sevoflurane general anesthesia. Pain Rep 6, e936 (2021).

25. Sprenger, T. et al. Imaging pain modulation by subanesthetic S-(+)-ketamine. Anesth. Analg. 103, 729–737 (2006).

26. Scheidegger, M. et al. Ketamine decreases resting state functional network connectivity in healthy subjects: implications for antidepressant drug action. PLoS One 7, e44799 (2012).

27. Bonhomme, V. et al. Resting-state Network-specific Breakdown of Functional Connectivity during Ketamine Alteration of Consciousness in Volunteers. Anesthesiology 125, 873–888 (2016).

28. Mueller, F. et al. Pharmacological fMRI: Effects of subanesthetic ketamine on resting-state functional connectivity in the default mode network, salience network, dorsal attention network and executive control network. Neuroimage Clin 19, 745–757 (2018).

29. Zacharias, N. et al. Ketamine effects on default mode network activity and vigilance: A randomized, placebo-controlled crossover simultaneous fMRI/EEG study. Hum. Brain Mapp. 41, 107–119 (2020).

30. Marguilho, M., Figueiredo, I. & Castro-Rodrigues, P. A unified model of ketamine’s dissociative and psychedelic properties. J. Psychopharmacol. 37, 14–32 (2023).

31. Kross, E., Berman, M. G., Mischel, W., Smith, E. E. & Wager, T. D. Social rejection shares somatosensory representations with physical pain. Proc. Natl. Acad. Sci. U. S. A. 108, 6270–6275 (2011).

32. Buhle, J. & Wager, T. D. Does meditation training lead to enduring changes in the anticipation and experience of pain? Pain 150, 382–383 (2010).

33. Wager, T. D. et al. Placebo-induced changes in FMRI in the anticipation and experience of pain. Science 303, 1162–1167 (2004).

34. Wager, T. D., Scott, D. J. & Zubieta, J.-K. Placebo effects on human mu-opioid activity during pain. Proc. Natl. Acad. Sci. U. S. A. 104, 11056–11061 (2007).

35. Wager, T. D. et al. An fMRI-based neurologic signature of physical pain. N. Engl. J. Med. 368, 1388–1397 (2013).

36. Wager, T. D. Using Neuroimaging to Understand Pain: Pattern Recognition and the Path from Brain Mapping to Mechanisms. The Brain Adapting with Pain: Contributions of Neuroimaging Technology to Pain Mechanisms 23–36 (2015).

37. Melzack, R. & Casey, K. L. Sensory, motivational, and central control determinants of pain: a new conceptual model. The skin senses (1968).

38. Tracey, I. & Mantyh, P. W. The cerebral signature for pain perception and its modulation. Neuron 55, 377–391 (2007).

39. Kenshalo, D. R., Jr & Isensee, O. Responses of primate SI cortical neurons to noxious stimuli. J. Neurophysiol. 50, 1479–1496 (1983).

40. Kenshalo, D. R., Jr, Chudler, E. H., Anton, F. & Dubner, R. SI nociceptive neurons participate in the encoding process by which monkeys perceive the intensity of noxious thermal stimulation. Brain Res. 454, 378–382 (1988).

41. Ploner, M., Freund, H. J. & Schnitzler, A. Pain affect without pain sensation in a patient with a postcentral lesion. Pain 81, 211–214 (1999).

42. Wiech, K. et al. Anterior insula integrates information about salience into perceptual decisions about pain. J. Neurosci. 30, 16324–16331 (2010).

43. Brown, C. A., Seymour, B., El-Deredy, W. & Jones, A. K. P. Confidence in beliefs about pain predicts expectancy effects on pain perception and anticipatory processing in right anterior insula. Pain 139, 324–332 (2008).

44. Rainville, P., Duncan, G. H., Price, D. D., Carrier, B. & Bushnell, M. C. Pain affect encoded in human anterior cingulate but not somatosensory cortex. Science 277, 968–971 (1997).

45. Tölle, T. R. et al. Region-specific encoding of sensory and affective components of pain in the human brain: a positron emission tomography correlation analysis. Ann. Neurol. 45, 40–47 (1999).

46. Rogers, R., Wise, R. G., Painter, D. J., Longe, S. E. & Tracey, I. An investigation to dissociate the analgesic and anesthetic properties of ketamine using functional magnetic resonance imaging. Anesthesiology 100, 292–301 (2004).

47. Denda, S. et al. Central nuclei and spinal pathways in feedback inhibitory spinal cord potentials in ketamine-anaesthetized rats. Br. J. Anaesth. 76, 258–265 (1996).

48. Ohtani, M. et al. Effects of ketamine on nociceptive cells in the medial medullary reticular formation of the cat. Anesthesiology 51, 414–417 (1979).

49. Thomson, A. M., West, D. C. & Lodge, D. An N-methylaspartate receptor-mediated synapse in rat cerebral cortex: a site of action of ketamine? Nature 313, 479–481 (1985).

50. Ren, K. Wind-up and the NMDA receptor: from animal studies to humans. Pain 59, 157–158 (1994).

51. Qu, X.-X. et al. Role of the spinal cord NR2B-containing NMDA receptors in the development of neuropathic pain. Exp. Neurol. 215, 298–307 (2009).

52. Eide, P. K. Wind-up and the NMDA receptor complex from a clinical perspective. Eur. J. Pain 4, 5–15 (2000).

53. Park, S. J., Chiang, C. Y., Hu, J. W. & Sessle, B. J. Neuroplasticity induced by tooth pulp stimulation in trigeminal subnucleus oralis involves NMDA receptor mechanisms. J. Neurophysiol. 85, 1836–1846 (2001).

54. Price, D. D., Mao, J., Frenk, H. & Mayer, D. J. The N-methyl-D-aspartate receptor antagonist dextromethorphan selectively reduces temporal summation of second pain in man. Pain 59, 165–174 (1994).

55. Larson, A. A. Interactions between ketamine and phencyclidine and dorsal root potentials (DRPs), evoked from the raphe nuclei. Neuropharmacology 23, 785–791 (1984).

56. Koizuka, S., Obata, H., Sasaki, M., Saito, S. & Goto, F. Systemic ketamine inhibits hypersensitivity after surgery via descending inhibitory pathways in rats. Can. J. Anaesth. 52, 498–505 (2005).

57. Zhu, X. et al. Evidence of a dissociation pattern in resting-state default mode network connectivity in first-episode, treatment-naive major depression patients. Biol. Psychiatry 71, 611–617 (2012).

58. Vesuna, S. et al. Deep posteromedial cortical rhythm in dissociation. Nature 586, 87–94 (2020).

59. Carhart-Harris, R. L. & Friston, K. J. REBUS and the Anarchic Brain: Toward a Unified Model of the Brain Action of Psychedelics. Pharmacol. Rev. 71, 316–344 (2019).

60. Satpute, A. B. & Lindquist, K. A. The Default Mode Network’s Role in Discrete Emotion. Trends Cogn. Sci. 23, 851–864 (2019).

61. Smallwood, J. et al. The default mode network in cognition: a topographical perspective. Nat. Rev. Neurosci. 22, 503–513 (2021).

62. Millière, R., Carhart-Harris, R. L., Roseman, L., Trautwein, F.-M. & Berkovich-Ohana, A. Psychedelics, Meditation, and Self-Consciousness. Front. Psychol. 9, 1475 (2018).

63. Letheby, C. & Gerrans, P. Self unbound: ego dissolution in psychedelic experience. Neurosci Conscious 2017, nix016 (2017).

64. Millière, R. Looking for the Self: Phenomenology, Neurophysiology and Philosophical Significance of Drug-induced Ego Dissolution. Front. Hum. Neurosci. 11, 245 (2017).

65. Wilkins, L. K., Girard, T. A. & Cheyne, J. A. Anomalous bodily-self experiences among recreational ketamine users. Cogn. Neuropsychiatry 17, 415–430 (2012).

66. Morgan, H. L. et al. Exploring the impact of ketamine on the experience of illusory body ownership. Biol. Psychiatry 69, 35–41 (2011).

67. Kaldewaij, R., et al. Ketamine reduces the neural distinction between self- and other-produced affective touch - a double-blind placebo-controlled study. PsyArXiv (2023) doi:10.31234/osf.io/w3ftk.

68. Vlisides, P. E. et al. Subanaesthetic ketamine and altered states of consciousness in humans. Br. J. Anaesth. 121, 249–259 (2018).

69. Gattuso, J. J. et al. Default Mode Network Modulation by Psychedelics: A Systematic Review. Int. J. Neuropsychopharmacol. 26, 155–188 (2023).

70. Mason, N. L. et al. Me, myself, bye: regional alterations in glutamate and the experience of ego dissolution with psilocybin. Neuropsychopharmacology 45, 2003–2011 (2020).

71. Carhart-Harris, R. L. et al. Neural correlates of the psychedelic state as determined by fMRI studies with psilocybin. Proc. Natl. Acad. Sci. U. S. A. 109, 2138–2143 (2012).

72. Carhart-Harris, R. L. et al. Neural correlates of the LSD experience revealed by multimodal neuroimaging. Proc. Natl. Acad. Sci. U. S. A. 113, 4853–4858 (2016).

73. Müller, F., Dolder, P. C., Schmidt, A., Liechti, M. E. & Borgwardt, S. Altered network hub connectivity after acute LSD administration. Neuroimage Clin 18, 694–701 (2018).

74. Siegel, J. S. et al. Psilocybin desynchronizes brain networks. medRxiv (2023) doi:10.1101/2023.08.22.23294131.

75. Madsen, M. K. et al. Psilocybin-induced changes in brain network integrity and segregation correlate with plasma psilocin level and psychedelic experience. Eur. Neuropsychopharmacol. 50, 121–132 (2021).

76. Palhano-Fontes, F. et al. The psychedelic state induced by ayahuasca modulates the activity and connectivity of the default mode network. PLoS One 10, e0118143 (2015).

77. Carhart-Harris, R. L. et al. The entropic brain: a theory of conscious states informed by neuroimaging research with psychedelic drugs. Front. Hum. Neurosci. 8, 20 (2014).

78. Kucyi, A. & Davis, K. D. The dynamic pain connectome. Trends Neurosci. 38, 86–95 (2015).

79. Zorn, J., Abdoun, O., Bouet, R. & Lutz, A. Mindfulness meditation is related to sensory-affective uncoupling of pain in trained novice and expert practitioners. Eur. J. Pain 24, 1301–1313 (2020).

80. Lutz, A., McFarlin, D. R., Perlman, D. M., Salomons, T. V. & Davidson, R. J. Altered anterior insula activation during anticipation and experience of painful stimuli in expert meditators. Neuroimage 64, 538–546 (2013).

81. Brown, C. A. & Jones, A. K. P. Meditation experience predicts less negative appraisal of pain: electrophysiological evidence for the involvement of anticipatory neural responses. Pain 150, 428–438 (2010).

82. Fan, C. et al. Effects of meditation on neural responses to pain: A systematic review and meta-analysis of fMRI studies. Neurosci. Biobehav. Rev. 162, 105735 (2024).

83. Cao, D. et al. Structure-based discovery of nonhallucinogenic psychedelic analogs. Science 375, 403–411 (2022).

84. Dunlap, L. E. et al. Identification of Psychoplastogenic N,N-Dimethylaminoisotryptamine (isoDMT) Analogues through Structure-Activity Relationship Studies. J. Med. Chem. 63, 1142–1155 (2020).

85. Cameron, L. P. et al. A non-hallucinogenic psychedelic analogue with therapeutic potential. Nature 589, 474–479 (2021).

86. Dong, C. et al. Psychedelic-inspired drug discovery using an engineered biosensor. Cell 184, 2779–2792.e18 (2021).

87. Lu, J. et al. An analog of psychedelics restores functional neural circuits disrupted by unpredictable stress. Mol. Psychiatry 26, 6237–6252 (2021).

88. Kaplan, A. L. et al. Bespoke library docking for 5-HT2A receptor agonists with antidepressant activity. Nature 610, 582–591 (2022).

89. Qu, Y., Chang, L., Ma, L., Wan, X. & Hashimoto, K. Rapid antidepressant-like effect of non-hallucinogenic psychedelic analog lisuride, but not hallucinogenic psychedelic DOI, in lipopolysaccharide-treated mice. Pharmacol. Biochem. Behav. 222, 173500 (2023).

90. Cunningham, M. J. et al. Pharmacological Mechanism of the Non-hallucinogenic 5-HT2A Agonist Ariadne and Analogs. ACS Chem. Neurosci. 14, 119–135 (2023).

91. Lewis, V. et al. A non-hallucinogenic LSD analog with therapeutic potential for mood disorders. Cell Rep. 42, 112203 (2023).

92. Lii, T. R. et al. Randomized trial of ketamine masked by surgical anesthesia in patients with depression. Nat Ment Health 1, 876–886 (2023).

93. Ballard, E. D. & Zarate, C. A., Jr. The role of dissociation in ketamine’s antidepressant effects. Nat. Commun. 11, 6431 (2020).

94. Nayak, S. M. et al. Control conditions in randomized trials of psychedelics: An ACTTION systematic review. J. Clin. Psychiatry 84, (2023).

95. Anand, A. et al. Attenuation of the neuropsychiatric effects of ketamine with lamotrigine: support for hyperglutamatergic effects of N-methyl-D-aspartate receptor antagonists. Arch. Gen. Psychiatry 57, 270–276 (2000).

96. Abdallah, C. G. et al. Prefrontal Connectivity and Glutamate Transmission: Relevance to Depression Pathophysiology and Ketamine Treatment. Biol Psychiatry Cogn Neurosci Neuroimaging 2, 566–574 (2017).

97. Deakin, J. F. W. et al. Glutamate and the neural basis of the subjective effects of ketamine: a pharmaco-magnetic resonance imaging study. Arch. Gen. Psychiatry 65, 154–164 (2008).

98. Doyle, O. M. et al. Quantifying the attenuation of the ketamine pharmacological magnetic resonance imaging response in humans: a validation using antipsychotic and glutamatergic agents. J. Pharmacol. Exp. Ther. 345, 151–160 (2013).

99. Woo, C.-W. et al. Separate neural representations for physical pain and social rejection. Nat. Commun. 5, 5380 (2014).

100. D’Souza, D. C. et al. Glycine transporter inhibitor attenuates the psychotomimetic effects of ketamine in healthy males: preliminary evidence. Neuropsychopharmacology 37, 1036–1046 (2012).

101. van Schalkwyk, G. I., Wilkinson, S. T., Davidson, L., Silverman, W. K. & Sanacora, G. Acute psychoactive effects of intravenous ketamine during treatment of mood disorders: Analysis of the Clinician Administered Dissociative State Scale. J. Affect. Disord. 227, 11–16 (2018).

102. Diazgranados, N. et al. A randomized add-on trial of an N-methyl-D-aspartate antagonist in treatment-resistant bipolar depression. Arch. Gen. Psychiatry 67, 793–802 (2010).

103. Murrough, J. W. et al. Ketamine for rapid reduction of suicidal ideation: a randomized controlled trial. Psychol. Med. 45, 3571–3580 (2015).

104. Singh, J. B. et al. A Double-Blind, Randomized, Placebo-Controlled, Dose-Frequency Study of Intravenous Ketamine in Patients With Treatment-Resistant Depression. Am. J. Psychiatry 173, 816–826 (2016).

105. Zarate, C. A., Jr & Niciu, M. J. Ketamine for depression: evidence, challenges and promise. World Psychiatry 14, 348–350 (2015).

106. Esteban, O. et al. fMRIPrep: a robust preprocessing pipeline for functional MRI. Nat. Methods 16, 111–116 (2019).

107. Siegel, J. S. et al. Statistical improvements in functional magnetic resonance imaging analyses produced by censoring high-motion data points. Hum. Brain Mapp. 35, 1981–1996 (2014).

108. Zunhammer, M., Bingel, U., Wager, T. D. & Placebo Imaging Consortium. Placebo Effects on the Neurologic Pain Signature: A Meta-analysis of Individual Participant Functional Magnetic Resonance Imaging Data. JAMA Neurol. 75, 1321–1330 (2018).

109. Han, X. et al. Effect sizes and test-retest reliability of the fMRI-based neurologic pain signature. Neuroimage 247, 118844 (2022).

110. López-Solà, M. et al. The neurologic pain signature responds to nonsteroidal anti-inflammatory treatment vs placebo in knee osteoarthritis. Pain Rep 7, e986 (2022).

111. Weber, K. A., Ii et al. Evidence for decreased Neurologic Pain Signature activation following thoracic spinal manipulation in healthy volunteers and participants with neck pain. Neuroimage Clin 24, 102042 (2019).

112. Whitfield-Gabrieli, S. & Nieto-Castanon, A. Conn: a functional connectivity toolbox for correlated and anticorrelated brain networks. Brain Connect. 2, 125–141 (2012).

113. Calhoun, V. D., Adali, T., Pearlson, G. D. & Pekar, J. J. A method for making group inferences from functional MRI data using independent component analysis. Hum. Brain Mapp. 14, 140–151 (2001).

114. Yeo, B. T. T. et al. The organization of the human cerebral cortex estimated by intrinsic functional connectivity. J. Neurophysiol. 106, 1125–1165 (2011).

115. Singmann, H. et al. afex: Analysis of factorial experiments. Comprehensive R Archive Network (CRAN). Preprint at (2016).

116. Barr, D. J., Levy, R., Scheepers, C. & Tily, H. J. Random effects structure for confirmatory hypothesis testing: Keep it maximal. J. Mem. Lang. 68, (2013).

