## Supplemental Materials for "The Analgesic and Dissociative Properties of Ketamine are Separate and Correspond to Distinct Neural Mechanisms"

Methods

Anatomical Preprocessing:

Within the FMRIPREP framework, each T1-weighted (T1w) image underwent intensity non-uniformity (INU) correction using 'N4BiasFieldCorrection’ v2.1.0 from 'AntsApplyTransforms' (ANTs version 2.2.0). The T1w reference was subsequently skull-stripped using the 'antsBrainExtraction.sh' workflow (from ANTs), with OASIS30-ANTs as the target template. Brain tissue segmentation (CSF, WM, GM) was conducted on the brain-extracted T1w using 'FAST' (FSL version 5.0.9). A T1w-reference map was generated after INU correction and registration of the T1w image using 'mri_robust_template' (FreeSurfer version 6.0.1). Volume-based spatial normalization to MNI152NLin2009cAsym standard space was accomplished through nonlinear registration with the 'antsRegistration' tool of ANTs version 2.2.0, employing brain-extracted versions of both the T1w reference and the T1w template, with the ICBM 152 nonlinear Asymmetrical template version 2009 as the target.

Functional Preprocessing:

A reference volume and its skull-stripped counterpart were generated using *fMRIPrep**^1^*, excluding susceptibility distortion correction. The BOLD reference was co-registered to the T1-weighted image using *MCFLIRT* (FSL 5.0.9) with nine degrees of freedom to account for residual distortions. Head-motion parameters were estimated relative to the BOLD reference. BOLD runs were slice-time corrected using AFNI’s 3dTshift (version 16.2.07), and their time series were resampled to native space after applying head motion correction transformations. Confounding time series—including framewise displacement (FD), DVARS, and three region-wise global signals (from CSF, white matter, and whole-brain masks)—were calculated. Physiological regressors were included using component-based noise correction (CompCor). After high-pass filtering the BOLD time series, both temporal (tCompCor) and anatomical (aCompCor) variants were computed. Six tCompCor components were extracted from the top 5% most variable voxels within the subcortical mask. For aCompCor, six components were extracted from the intersection of the subcortical mask with the CSF and WM masks (projected to native space). For each decomposition, only the k components explaining at least 50% of the variance were retained. Given the importance of controlling for physiological noise in ketamine studies^2^, we used the PhysIO toolbox^3^; <https://translationalneuromodeling.github.io/tapas>) to model respiratory, cardiac, and vascular activity and incorporated the output into the CompCor regressors. All resampling was performed using a single interpolation step that combined relevant transformations. Volumetric resampling was performed using ANTs with Lanczos interpolation to minimize smoothing artifacts. Surface resampling was performed using FreeSurfer’s mri_vol2surf. Many internal operations of *fMRIPrep* employed *Nilearn*, particularly for BOLD processing (see: <https://fmriprep.readthedocs.io/en/stable/workflows.html>). In first-level analyses, the confounds file included the following regressors: six head motion estimates, their quadratic terms and temporal derivatives (24 total), the standard deviation of DVARS, six aCompCor components, framewise displacement (FD), and 18 physiological regressors. Volumes with FD > 0.4 mm were flagged as motion outliers and scrubbed from analysis (Siegel et al., 2014). Finally, spatial smoothing was applied using SPM12 with a 6 mm full-width at half-maximum kernel.

Network extraction procedure

Group-level ICA was conducted using the CONN toolbox, applying the G1 FastICA algorithm as described previously^4^. The number of components was fixed at 20. A three-step principal component analysis (PCA) was applied during preprocessing. This process initially reduced individual participant data, segmented it into distinct independent components, and subsequently estimated subject-specific spatial maps and corresponding time courses using the GICA3 algorithm. Following ICA decomposition, a nonparametric analysis was applied to the group-level spatial component maps to assess statistical significance across subjects.

To label the extracted components, we employed CONN’s built-in network labeling tool, which quantifies the spatial overlap between each independent component and the canonical network parcellation developed by Yeo et al^5^. Based on this overlap, five components were identified as having meaningful correspondence to large-scale brain networks. These components were labeled as , DMN, SN, FPN, SMN, DAN, Visual.

Connectivity Analysis

Connectivity analysis was performed using a predefined set of anatomically defined ROIs corresponding to each of the identified functional networks. For each session and participant, a mean BOLD time course was extracted across selected voxels within each ROI. Pairwise Pearson correlation coefficients were then computed between all ROI pairs within each network using bivariate regression models. These coefficients were transformed into z-scores using Fisher’s r-to-z transformation to facilitate group-level statistical analyses. The primary contrast of interest in the second-level ROI-to-ROI analysis focused on detecting changes in within-network connectivity between ketamine and placebo sessions. Within-network connectivity was defined as the mean correlation between all ROI pairs belonging to the same network.

|  | **Coordinates (MNI)** | | | **Network** |
| --- | --- | --- | --- | --- |
| **ROI Label** | X | Y | Z |  |
| Posterior Cingulate Cortex (PCC) | 0 | -54 | 31 | DMN |
| Inferior Parietal Lobule R | 44 | -65 | 33 | DMN |
| Inferior Parietal Lobule L | -37 | 73 | 38 | DMN |
| Ventromedial Prefrontal Cortex | 0 | 54 | -6 | DMN |
| Parahippocampal Gyrus R | 25 | -36 | -17 | DMN |
| Parahippocampal Gyrus L | -27 | -40 | -13 | DMN |
| Anterior Cingulate Cortex | 0 | 18 | 41 | SN |
| Dorsolateral Prefrontal Cortex R | 32 | 45 | 28 | SN |
| Dorsolateral Prefrontal Cortex L | -33 | 44 | 27 | SN |
| Supramarginal Gyrus R | 53 | -43 | 40 | SN |
| Supramarginal Gyrus L | -36 | -50 | 43 | SN |
| Insular Cortex R | 45 | 16 | 0 | SN |
| Insular Cortex L | -40 | 15 | 3 | SN |
| Posterior Cingulate Cortex | 0 | -25 | 28 | SN |
| Mid Temporal Lobe L | -56 | -41 | 9 | Frontoparietal |
| Middle Frontal Gyrus R | 52 | 24 | 4 | Frontoparietal |
| Middle Frontal Gyrus L | -51 | 22 | 2 | Frontoparietal |
| Precuneus R | 2 | -55 | 43 | Frontoparietal |
| Precentral Gyrus L | -43 | 1 | 53 | Frontoparietal |
| Parietal Supplementary Area | -22 | -79 | 37 | Frontoparietal |
| Somatosensory Area R | 50 | -12 | 33 | Sensorimotor |
| Somatosensory Area L | -52 | -12 | 28 | Sensorimotor |
| Sensorymotor Cortex | 2 | -23 | 60 | Sensorimotor |
| Intraprietal Sulcus R | 44 | -32 | 52 | Dorsal Attention |
| Intraprietal Sulcus L | -39 | -28 | 53 | Dorsal Attention |
| Supplementary Motor Area | -2 | -4 | 53 | Dorsal Attention |
| Premotor Area L | -57 | 4 | 36 | Dorsal Attention |
| Premotor Area R | 58 | 9 | 33 | Dorsal Attention |
| Occipital lobe R | 25 | -65 | 30 | Visual |
| Occipital lobe L | -18 | -70 | -4 | Visual |

**Supplemental Table S1.** Regions of interest (ROIs) used in the within-network connectivity analysis, grouped by functional network affiliation. Network labels correspond to components derived from group ICA and matched to canonical functional networks as defined by Yeo et al. ^5^. These ROIs served as the basis for calculating within-network connectivity metrics in the ROI-to-ROI analysis.

Results

Effect of ketamine on dissociative state

| **Effect** | **df** | **F** | **p-value** |
| --- | --- | --- | --- |
| Time Point | 2, 476.15 | 104.25 | <.001 |
| Session | 1, 32.07 | 57.44 | <.001 |
| Subscale | 2, 464.23 | 64.1 | <.001 |
| Time Point X Session | 2, 477.73 | 109.51 | <.001 |
| Time Point X Subscale | 4, 464.23 | 8.19 | <.001 |
| Session X Subscale | 2, 464.23 | 9.87 | <.001 |
| Time Point X Session X Subscale | 4, 464.23 | 11.15 | <.001 |

**Supplemental Table S2.** Output of the Linear mixed-effects model: rating~time_point*session*sub_scale+(1+session|subji). This model examines the effects of time point (pre-infusion, post-bolus, end of infusion), session (ketamine vs. placebo), and their interaction on CADSS dissociation ratings, including random intercepts and slopes for session by subject.

| **Contrast** | **Time Point** | **Estimate** | **SE** | **df** | **t-ratio** | **p-value** |
| --- | --- | --- | --- | --- | --- | --- |
| Ketamine–Placebo | End of infusion | 3.21 | 0.555 | 97.9 | 5.785 | <.0001 |
| Ketamine–Placebo | Post-bolus | 7.13 | 0.509 | 76.4 | 13.999 | <.0001 |
| Ketamine–Placebo | Pre-infusion | -1.05 | 0.519 | 80.6 | -2.019 | 0.0468 |

**Supplemental Table S3.** Post-Hoc tests for each time point in the placebo - ketamine contrast for Supplemental Table S2.

| **Contrast** | **Session** | **Subscale** | **Estimate** | **SE** | **df** | **t-ratio** | **p-value** |
| --- | --- | --- | --- | --- | --- | --- | --- |
| Post-bolus–End_of_infusion | Placebo | Amnesia | -0.041 | 0.714 | 471 | -0.057 | .955 |
| Pre-infusion–Post-bolus | Placebo | Amnesia | -0.233 | 0.678 | 466 | -0.344 | .952 |
| Post-bolus–End_of_infusion | Ketamine | Amnesia | 2.431 | 0.707 | 468 | 3.439 | .001 |
| Pre-infusion–Post-bolus | Ketamine | Amnesia | -4.556 | 0.672 | 465 | -6.777 | <.001 |
| Post-bolus–End_of_infusion | Placebo | Depersonalization | 0.185 | 0.714 | 471 | 0.260 | .952 |
| Pre-infusion–Post-bolus | Placebo | Depersonalization | 0.109 | 0.678 | 466 | 0.161 | .952 |
| Post-bolus–End_of_infusion | Ketamine | Depersonalization | 3.151 | 0.707 | 468 | 4.458 | <.001 |
| Pre-infusion–Post-bolus | Ketamine | Depersonalization | -7.011 | 0.672 | 465 | -10.429 | <.001 |
| Post-bolus–End_of_infusion | Placebo | Derealisation | 0.118 | 0.714 | 471 | 0.165 | .952 |
| Pre-infusion–Post-bolus | Placebo | Derealisation | 0.425 | 0.678 | 466 | 0.626 | .911 |
| Post-bolus–End_of_infusion | Ketamine | Derealisation | 6.448 | 0.707 | 468 | 9.121 | <.001 |
| Pre-infusion–Post-bolus | Ketamine | Derealisation | -12.667 | 0.672 | 465 | -18.843 | <.001 |

**Supplemental Table S4.** Post-Hoc tests for each time point in the placebo - ketamine and time point contrast for Supplemental Table S2.

The effects of Ketamine on pain related brain activation

| **Effect** | **df** | **F** | **p-value** |
| --- | --- | --- | --- |
| Intensity | 1, 702 | 124.27 | < 0.001 |
| Session | 1, 702 | 43.05 | < 0.001 |
| ROI | 6, 702 | 8.11 | < 0.001 |
| Intensity × Session | 1, 702 | 16.45 | < 0.001 |
| Intensity × ROI | 6, 702 | 0.47 | 0.831 |
| Session × ROI | 6, 702 | 0.17 | 0.986 |
| Intensity × Session × ROI | 6, 702 | 0.06 | 0.999 |

**Supplemental Table S5*:*** *Full model output for The effects of Ketamine on pain related brain activation. beta~intensity*session*ROI+(1|subji).*

| **ROI** | **Contrast** | **Estimate** | **SE** | **df** | **t** | **p-value** |
| --- | --- | --- | --- | --- | --- | --- |
| Ant. Insula (L) | Placebo- Ketamine | 0.335 | 0.136 | 338 | 2.463 | 0.014 |
| Ant. Insula (R) | Placebo- Ketamine | 0.401 | 0.136 | 338 | 2.953 | 0.003 |
| dACC | Placebo- Ketamine | 0.403 | 0.136 | 338 | 2.965 | 0.003 |
| DLPFC (R) | Placebo- Ketamine | 0.390 | 0.136 | 338 | 2.868 | 0.004 |
| S1 (R) | Placebo- Ketamine | 0.548 | 0.136 | 338 | 4.030 | < 0.001 |
| S2 (L) | Placebo- Ketamine | 0.408 | 0.136 | 338 | 3.000 | 0.003 |
| S2 (R) | Placebo- Ketamine | 0.458 | 0.136 | 338 | 3.367 | < 0.001 |

**Supplemental Table S6*:*** *Post-Hoc tests for each ROI in the placebo - ketamine contrast*

| **ROI** | **Session** | **Spearman’s ρ** | **p-value** |
| --- | --- | --- | --- |
| Ant. Insula (L) | Placebo | 0.528 | 0.005 |
| Ant. Insula (L) | Ketamine | 0.656 | < 0.001 |
| Ant. Insula (R) | Placebo | 0.463 | 0.015 |
| Ant. Insula (R) | Ketamine | 0.561 | 0.002 |
| DLPFC (R) | Placebo | 0.213 | 0.286 |
| DLPFC (R) | Ketamine | 0.570 | 0.002 |
| dACC | Placebo | 0.359 | 0.066 |
| dACC | Ketamine | 0.651 | < 0.001 |
| S1 (R) | Placebo | 0.134 | 0.504 |
| S1 (R) | Ketamine | 0.448 | 0.019 |
| S2 (L) | Placebo | 0.460 | 0.016 |
| S2 (L) | Ketamine | 0.344 | 0.079 |
| S2 (R) | Placebo | 0.427 | 0.026 |
| S2 (R) | Ketamine | 0.491 | 0.009 |

**Supplemental Table S7*:*** Spearman correlation between ROI activation difference and pain rating difference. This table reports Spearman’s rank correlation coefficients (ρ) and associated p-values for the relationship between activation difference (high – low pain) in each region of interest (ROI) and the corresponding subjective pain rating difference, separately for the placebo and ketamine sessions.

| **ROI** | **Pain‑Only p** | **Depersonalization p** | **Derealization p** | **Amnesia p** | **Model Comparison p** |
| --- | --- | --- | --- | --- | --- |
| Ant. Insula (L) | 0.0009 | 0.825 | 0.98 | 0.827 | 0.969 |
| Ant. Insula (R) | 0.0030 | 0.58 | 0.7 | 0.703 | 0.853 |
| dACC | 0.0030 | 0.892 | 0.965 | 0.655 | 0.922 |
| DLPFC (R) | 0.0054 | 0.62 | 0.661 | 0.449 | 0.761 |
| S1 (R) | 0.0885 | 0.384 | 0.875 | 0.258 | 0.632 |
| S2 (L) | 0.0400 | 0.6 | 0.312 | 0.795 | 0.777 |
| S2 (R) | 0.0330 | 0.983 | 0.662 | 0.845 | 0.968 |

| **ROI** | **F** | **df1** | **df2** | **Adj R² (Pain‑Only)** | **Adj R² (Full)** | **AIC (Pain‑Only)** | **AIC (Full)** |
| --- | --- | --- | --- | --- | --- | --- | --- |
| Ant. Insula (L) | 0.081 | 3 | 22 | 0.429 | 0.358 | 44.6 | 50.3 |
| Ant. Insula (R) | 0.261 | 3 | 22 | 0.394 | 0.335 | 50.8 | 55.9 |
| dACC | 0.160 | 3 | 22 | 0.375 | 0.305 | 55.9 | 61.3 |
| DLPFC (R) | 0.390 | 3 | 22 | 0.336 | 0.283 | 50.1 | 54.7 |
| S1 (R) | 0.584 | 3 | 22 | 0.199 | 0.156 | 53.6 | 57.6 |
| S2 (L) | 0.367 | 3 | 22 | 0.173 | 0.105 | 59.5 | 64.2 |
| S2 (R) | 0.084 | 3 | 22 | 0.208 | 0.110 | 53.8 | 59.5 |

**Supplemental Table S8.** *Model results for pain-related brain activation during ketamine administration.* This table presents results from two linear models fitted separately for each region of interest (ROI). The *Pain Only Model* includes subjective pain ratings as the sole predictor of brain activation. The *Pain And Dissociation Model* adds dissociation subscale scores (Depersonalization, Derealization, Amnesia) to the model. The table reports p-values for each dissociation predictor, the model comparison p-value (*P Model Comparison*) based on ANOVA, adjusted R² for both models, and model AICs. A significant p-value in the model comparison supports the added explanatory value of including dissociation scores.

| **Outcome** | **Pain Only Model p** | **Depersonalization p** | **Derealization p** | **Amnesia p** | **ANOVA p** | **F-stat** | **df1** | **df2** |
| --- | --- | --- | --- | --- | --- | --- | --- | --- |
| NPS | 0.01 | 0.568 | 0.067 | 0.717 | 0.277 | 1.37 | 3 | 22 |

| **Adj. R² (Pain Only Model Model)** | **Adj. R² (Pain And Dissociation Model)** | **AIC (Pain Only Model )** | **AIC (Pain And Dissociation Model)** |
| --- | --- | --- | --- |
| 0.27 | 0.3018 | 244 | 245 |

**Supplemental Table S9.** *Model results for pain-related multivariate brain activation during ketamine administration.* This table presents results from two linear models fitted separately predicting the multi-voxel biomarker for the subjective intensity of pain: the Neurologic Pain Signature . The *Pain Only Model* includes subjective pain ratings as the sole predictor of brain activation. The *Pain And Dissociation Model* adds dissociation subscale scores (Depersonalization, Derealization, Amnesia) to the model. The table reports p-values for each dissociation predictor, the model comparison p-value (*P Model Comparison*) based on ANOVA, adjusted R² for both models, and model AICs.

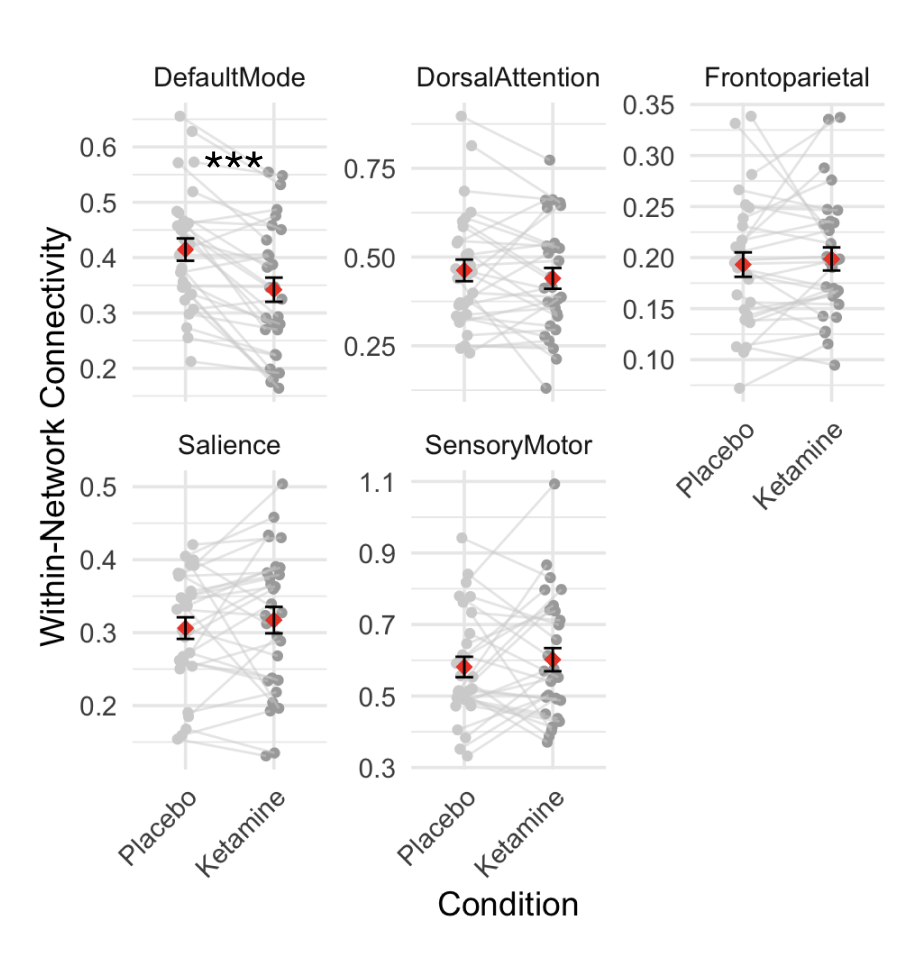

**Figure S.1.** *Effects of ketamine on within-network functional connectivity across brain networks.* Connectivity values were extracted from the pain task fMRI data in each session and computed within the following networks: Default Mode Network, Dorsal Attention Network, Frontoparietal Network, Salience Network, and Somato Motor Network . Individual subject trajectories are shown in grey, with red diamonds representing the group mean and error bars indicating the standard error of the mean. **** p<.001.*

References

1.	Esteban, O. *et al.* fMRIPrep: a robust preprocessing pipeline for functional MRI. *Nat. Methods* **16**, 111–116 (2019).

2.	McMillan, R. & Muthukumaraswamy, S. D. The neurophysiology of ketamine: an integrative review. *Rev. Neurosci.* **31**, 457–503 (2020).

3.	Kasper, L. *et al.* The PhysIO Toolbox for Modeling Physiological Noise in fMRI Data. *J. Neurosci. Methods* **276**, 56–72 (2017).

4.	Calhoun, V. D., Adali, T., Pearlson, G. D. & Pekar, J. J. A method for making group inferences from functional MRI data using independent component analysis. *Hum. Brain Mapp.* **14**, 140–151 (2001).

5.	Yeo, B. T. T. *et al.* The organization of the human cerebral cortex estimated by intrinsic functional connectivity. *J. Neurophysiol.* **106**, 1125–1165 (2011).
